# Conserved heavy/light contacts and germline preferences revealed by a large-scale analysis of natively paired human antibody sequences and structural data

**DOI:** 10.1101/2024.12.20.629642

**Authors:** Pawel Dudzic, Dawid Chomicz, Weronika Bielska, Igor Jaszczyszyn, Michał Zieliński, Bartosz Janusz, Sonia Wróbel, Marguerite-Marie Le Pannérer, Andrew Philips, Prabakaran Ponraj, Sandeep Kumar, Konrad Krawczyk

## Abstract

Antibody next-generation sequencing (NGS) datasets have become crucial to develop computational models addressing this successful class of therapeutics. Although antibodies are composed of both heavy and light chains, most NGS sequencing depositions provide them in unpaired form, reducing their utility. Here we introduce PairedAbNGS, a novel database with paired heavy/light antibody chains. To the best of our knowledge, this is the largest resource for paired natural antibody sequences with 58 bioprojects and over 14 million assembled productive sequences. We make the database accessible at https://naturalantibody.com/paired-ab-ngs as a valuable tool for biological and machine-learning applications. Using this dataset, we investigated heavy and light chain variable (V) gene pairing preferences and found significant biases beyond gene usage frequencies, possibly due to receptor editing favoring less autoreactive combinations. Analyzing the available antibody structures from the Protein Data Bank, we studied conserved contact residues between heavy and light chains, particularly interactions between the CDR3 region of one chain and the FWR2 region of the opposite chain. Examination of amino acid pairs at key contact sites revealed significant deviations of amino acids distributions compared to random pairings, in the heavy chain’s CDR3 region contacting the opposite chain, indicating specific interactions might be crucial for proper chain pairing. This observation is further reinforced by preferential IGHV-IGLJ and IGLV-IGHJ pairing preferences. We hope that both our resources and the findings would contribute to improving the engineering of biological drugs.

## Introduction

Therapeutic antibodies are rapidly emerging as the most successful class of biotherapeutics, with their yearly number of regulatory approvals reaching the same levels as the small molecule drugs (Senior, 2024). However, the initial generation of antibodies against a given antigen primarily requires the use of experimental methods, such as animal immunization, hybridomas, or phage/yeast display libraries (Gray *et al*., 2020).

In addition to being costly and time-consuming, experimental methods only work well for antigens that are ‘well-behaved’ under laboratory conditions (Stephens and Wilkinson, 2024). That is, their expression yields are high, in addition to them being conformationally stable and highly soluble in vitro. In the case of challenging targets such as GPCRs, claudins and other membrane-associated proteins that demonstrate conformational instability and/or low solubility outside of their cellular milieu, the task of producing sufficient amounts of antigens to initiate experimental generation of antibody binders becomes daunting.

To overcome these limitations and to expand the antigen space druggable by biotherapeutics to include the antigens where the experimental approaches may not work well, it is crucial to explore novel routes of antibody generation. Computation can be very helpful in this regard. This is because the availability of large antibody repertoires via next-generation sequencing (NGS) has coincided with the immense success of methods in machine learning theory and generalizations (Vidyasagar, 1996; Wilman *et al*., 2022). Drawing from publicly available antibody sequencing data either by data mining or machine learning modeling, it is now feasible to devise small and focused antigen-specific as well as larger antigen-agnostic antibody libraries (Bauer *et al*., 2023; Porebski *et al*., 2024). However, this endeavor requires careful construction and curation of underlying antibody sequences for deep learning, along with benchmarking different generative AI algorithms for their suitability towards specific biological purposes.

Paired antibody sequence databases are crucial for advancing our understanding of how immune responses are initiated and developed in the human body when challenged by an antigen. A crucial question is as follows: Development of IgG antibodies against an antigen requires class switching (IgM → IgG) and affinity maturation (Stavnezer and Schrader, 2014). Affinity maturation is often thought to be limited to V(D)J recombination and somatic hypermutation (Mishra and Mariuzza, 2018). However, does it also require selection of specific germline pairing between the light and heavy chains (Enzelberger *et al*., 2016)?. This question has been so far neglected because of lack of data available for naive B-cell and immunized antibody repertoires. Understanding this question is crucial for developing therapeutic antibodies, and improving reagent and diagnostic antibodies. However, existing databases face limitations in diversity, data quality, and accessibility. The original version of OAS only contained unpaired reads (Kovaltsuk *et al*., 2018). Subsequent update collated ca. 120k paired sequences from five studies. The resource is regularly updated, however certain studies such as one by Jaffe et al. account for most of its diversity (Jaffe *et al*., 2022). The presence of memory cells and the phenomenon of functional coherence can further limit the overall diversity represented in the dataset. We have recently demonstrated that by automatic mining of public repositories, it is possible to identify a large number of antibody-containing depositions (Dudzic, Chomicz, *et al*., 2024). Though at the time, because of the technical simplicity of processing unpaired reads and the corresponding complexity of doing so with the paired datasets, we only made unpaired data available.

To address the lack of diverse paired antibody data, we created a novel database of human and mouse-paired antibody sequences (PairedAbNGS) derived from publicly available data in the NCBI Sequence Read Archive (SRA), focusing specifically on projects utilizing high throughput single cell analyses via the 10x Chromium 5’ pipeline. Single-cell technology has revolutionized our ability to capture and analyze individual cells, providing unprecedented insights into cellular heterogeneity and function (Pearson *et al*., 2022). In the context of antibody research, single-cell RNA sequencing (scRNA-seq) enables the simultaneous capture of paired heavy and light chain sequences from individual B-cells, overcoming the limitations of bulk sequencing approaches (Zhang *et al*., 2021). Previously, barcoded technology microfluidic methods for single-B-cell isolation led to the identification of millions of paired human antibody sequences, as demonstrated in studies by Rajan et al. (Rajan *et al*., 2018) and Wang et al. (Wang *et al*., 2018). However, the actual number of unique sequences recovered may be influenced by factors such as the diversity of the B-cell population being studied, the efficiency of the isolation and sequencing processes, and the depth of sequencing coverage. The 10x Chromium 5’ pipeline, in particular, offers high-throughput single-cell analysis with improved full-length transcript coverage, making it an ideal platform for generating paired antibody sequences. By leveraging this technology and curating data from multiple studies, our database aims to provide the clearest landscape to date on the natural pairing of heavy and light chains among antibodies.

It is known that heavy/light preferences affect the binding of the antibodies to antigens (Dondelinger *et al*., 2018; Fernández-Quintero *et al*., 2020), with multiple computational studies on the smaller datasets. Via such studies certain genetic preferences, as well as key residues on the interface affecting the Vh/Vl conformation were proposed (Abhinandan and Martin, 2010; Dunbar *et al*., 2013; Bujotzek *et al*., 2015, 2016; Boron and Martin, 2023). In addition, heavy and light chain pairings belonging to IGHV3 and IGKV1 genetic loci are most frequent among the marketed antibody-based biotherapeutics.

Leveraging the largest paired antibody collection to date, we have studied the pairing preferences at the gene and individual residue levels. We explored inter-chain interaction patterns where we observed that the CDR3 of each chain is in contact with the FWR2 of the opposing chain. We hope that our analysis and novel dataset will act as a good basis for future studies of pairing preferences and will aid in antibody engineering.

## Results

### Overview of database curation

NCBI SRA served as the primary source of sequencing data, with metadata imported from EBI ENA in May 2024. We focused on projects containing sequencing experiments labeled as “TRANSCRIPTOMIC SINGLE CELL’’ and used a large language model (Gemini) to identify antibody-related studies (Gemini Team *et al*., 2023). This process involved creating a specific prompt for the Gemma language model (Gemma Team *et al*., 2024) and manually verifying the results. Only projects using the 10x Chromium 5’ library preparation and focusing on human or mouse data were included. We identified 58 suitable studies containing 2482 sequencing experiments, resulting in over 14 million assembled productive sequences. Reads were processed using sra-toolkit and TRUST4, with annotation performed using the RIOT (Dudzic, Janusz, *et al*., 2024) program with OGRDB (Lees *et al*., 2020) as a primary source of gene references. The extracted data were divided by barcodes, and only productive (without the stop codon in translation and frameshift between V and J gene segments) chains were retained as annotated by RIOT flag “productive”. Sequences were sorted by coverage, and only the top heavy and light chain pair for each barcode was kept. This filtering and processing ensured high-quality data for further analysis. Details of the procedure are laid out in the Methods section.

### Database statistics

The database consists of 58 studies with 2482 single cell sequencing experiments totaling 14,401,268 productive assembled nucleotide sequences. This yields 7,200,634 million amino acid sequences of which 4,280,331 are unique and span all three complementarity-determining regions (CDRs) along with framework regions (FRs) in both heavy and light chains. The cumulative number of antibody-focused single cell projects deposited in SRA (Sayers *et al*., 2022) presents notable growth (Figure 1). Project identification was performed in May 2024. The high rate of growth in the number of bioprojects speaks in favor of an automated identification pipeline, as developed in this and our previous study (Dudzic, Chomicz, *et al*., 2024).

**Figure 1.**
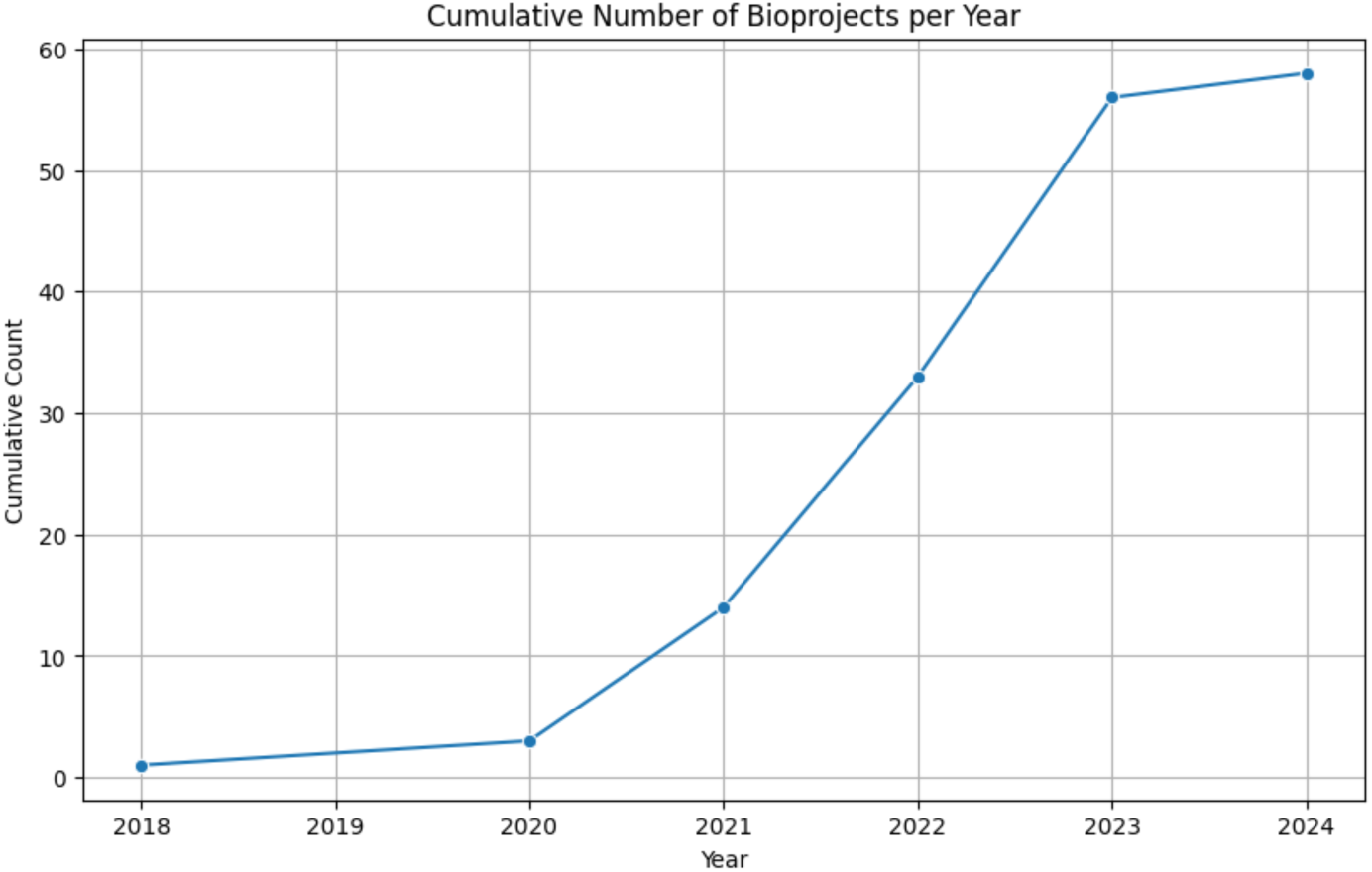
Cumulative number of single-cell (paired heavy/light) bioprojects published per year. Please note since the identification was performed in May 2024, the numbers for 2024 are incomplete and are therefore the increment is lower than for previous years.

The overall ratio of lambda and kappa chains in human samples closely aligns with the established consensus ratios: 60% kappa and 40% lambda chains as shown in Table 1 (Collins and Jackson, 2018). However, project-specific variations are evident in Table 2, which details the proportions for each individual project. In rodents, there is a significant bias toward kappa chains, with observed proportions being 95% kappa and 5% lambda. Interestingly, our dataset reveals that the overall kappa to lambda ratio in mice is 90% kappa and 10% lambda (Table 1), which is not as expected. This bias could be attributed to the PRJNA1082281 project, which focuses on extrafollicular plasmablasts and where identified lambda chains constitute 33% of all light sequences. A closer examination of the mouse projects in Table 4 shows that most adhere to the canonical 95:5 kappa to lambda ratio. Grouped metrics of chain proportions, such as mean and standard deviation are presented in Table 3 and Table 5 for human and mouse projects respectively.

**Table 1.** Number of heavy/light chain sequences and Light chain proportions for each organism.

| organism | igh | igk | igl | igk_ratio | igl_ratio |
| --- | --- | --- | --- | --- | --- |
| human | 6566748 | 3959197 | 2607551 | 0.603 | 0.397 |
| mouse | 633886 | 570298 | 63588 | 0.9 | 0.1 |

**Table 2.**
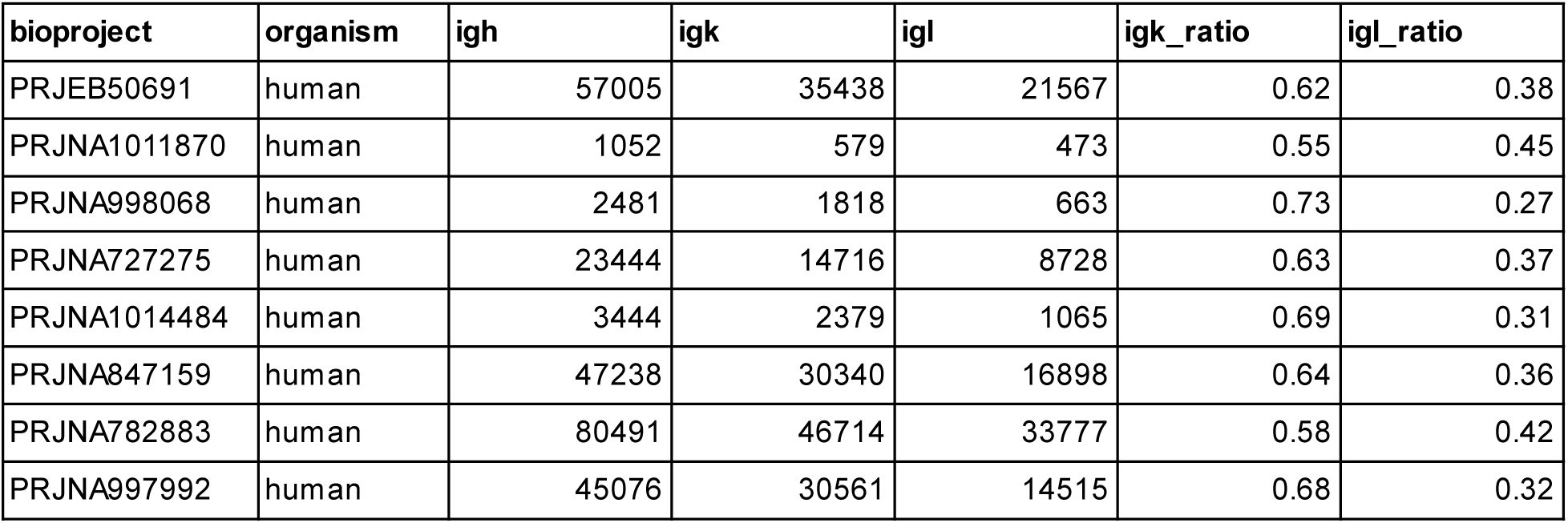
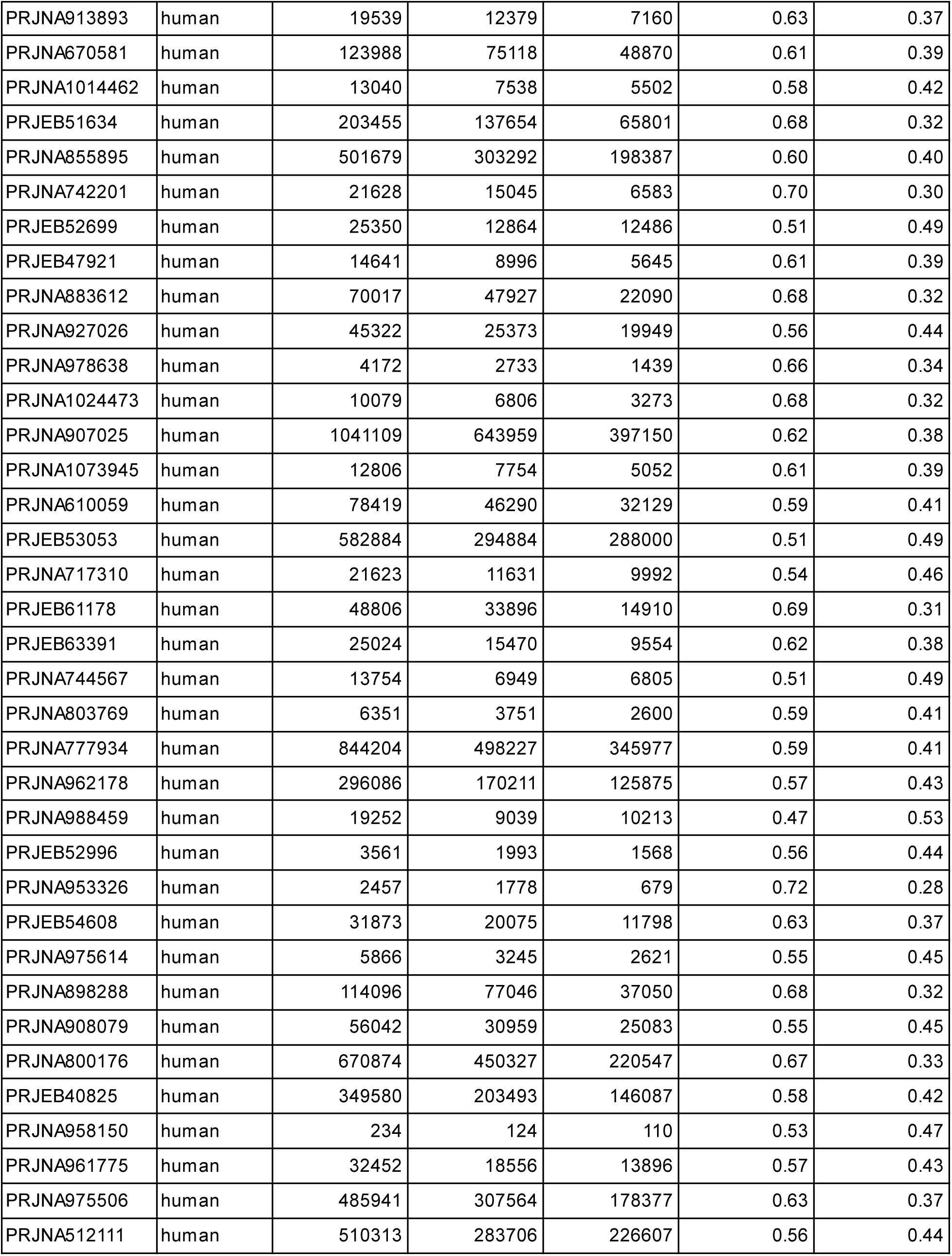
Number of heavy /light chain sequences and Light chain proportions for each human project.

| bioproject | organism | igh | igk | igl | igk_ratio | igl_ratio |
| --- | --- | --- | --- | --- | --- | --- |
| PRJEB50691 | human | 57005 | 35438 | 21567 | 0.62 | 0.38 |
| PRJNA1011870 | human | 1052 | 579 | 473 | 0.55 | 0.45 |
| PRJNA998068 | human | 2481 | 1818 | 663 | 0.73 | 0.27 |
| PRJNA727275 | human | 23444 | 14716 | 8728 | 0.63 | 0.37 |
| PRJNA1014484 | human | 3444 | 2379 | 1065 | 0.69 | 0.31 |
| PRJNA847159 | human | 47238 | 30340 | 16898 | 0.64 | 0.36 |
| PRJNA782883 | human | 80491 | 46714 | 33777 | 0.58 | 0.42 |
| PRJNA997992 | human | 45076 | 30561 | 14515 | 0.68 | 0.32 |
| PRJNA913893 | human | 19539 | 12379 | 7160 | 0.63 | 0.37 |
| PRJNA670581 | human | 123988 | 75118 | 48870 | 0.61 | 0.39 |
| PRJNA1014462 | human | 13040 | 7538 | 5502 | 0.58 | 0.42 |
| PRJEB51634 | human | 203455 | 137654 | 65801 | 0.68 | 0.32 |
| PRJNA855895 | human | 501679 | 303292 | 198387 | 0.60 | 0.40 |
| PRJNA742201 | human | 21628 | 15045 | 6583 | 0.70 | 0.30 |
| PRJEB52699 | human | 25350 | 12864 | 12486 | 0.51 | 0.49 |
| PRJEB47921 | human | 14641 | 8996 | 5645 | 0.61 | 0.39 |
| PRJNA883612 | human | 70017 | 47927 | 22090 | 0.68 | 0.32 |
| PRJNA927026 | human | 45322 | 25373 | 19949 | 0.56 | 0.44 |
| PRJNA978638 | human | 4172 | 2733 | 1439 | 0.66 | 0.34 |
| PRJNA1024473 | human | 10079 | 6806 | 3273 | 0.68 | 0.32 |
| PRJNA907025 | human | 1041109 | 643959 | 397150 | 0.62 | 0.38 |
| PRJNA1073945 | human | 12806 | 7754 | 5052 | 0.61 | 0.39 |
| PRJNA610059 | human | 78419 | 46290 | 32129 | 0.59 | 0.41 |
| PRJEB53053 | human | 582884 | 294884 | 288000 | 0.51 | 0.49 |
| PRJNA717310 | human | 21623 | 11631 | 9992 | 0.54 | 0.46 |
| PRJEB61178 | human | 48806 | 33896 | 14910 | 0.69 | 0.31 |
| PRJEB63391 | human | 25024 | 15470 | 9554 | 0.62 | 0.38 |
| PRJNA744567 | human | 13754 | 6949 | 6805 | 0.51 | 0.49 |
| PRJNA803769 | human | 6351 | 3751 | 2600 | 0.59 | 0.41 |
| PRJNA777934 | human | 844204 | 498227 | 345977 | 0.59 | 0.41 |
| PRJNA962178 | human | 296086 | 170211 | 125875 | 0.57 | 0.43 |
| PRJNA988459 | human | 19252 | 9039 | 10213 | 0.47 | 0.53 |
| PRJEB52996 | human | 3561 | 1993 | 1568 | 0.56 | 0.44 |
| PRJNA953326 | human | 2457 | 1778 | 679 | 0.72 | 0.28 |
| PRJEB54608 | human | 31873 | 20075 | 11798 | 0.63 | 0.37 |
| PRJNA975614 | human | 5866 | 3245 | 2621 | 0.55 | 0.45 |
| PRJNA898288 | human | 114096 | 77046 | 37050 | 0.68 | 0.32 |
| PRJNA908079 | human | 56042 | 30959 | 25083 | 0.55 | 0.45 |
| PRJNA800176 | human | 670874 | 450327 | 220547 | 0.67 | 0.33 |
| PRJEB40825 | human | 349580 | 203493 | 146087 | 0.58 | 0.42 |
| PRJNA958150 | human | 234 | 124 | 110 | 0.53 | 0.47 |
| PRJNA961775 | human | 32452 | 18556 | 13896 | 0.57 | 0.43 |
| PRJNA975506 | human | 485941 | 307564 | 178377 | 0.63 | 0.37 |
| PRJNA512111 | human | 510313 | 283706 | 226607 | 0.56 | 0.44 |

**Table 3.** Grouped metrics of chain proportions of sequences from human projects.

| metric | igh | igk | igl | igk_ratio | igl_ratio |
| --- | --- | --- | --- | --- | --- |
| mean | 149244,27 | 89981,75 | 59262,52 | 0,61 | 0,39 |
| std | 249344,59 | 150902,55 | 99799,53 | 0,06 | 0,06 |
| min | 234,00 | 124,00 | 110,00 | 0,06 | 0,27 |
| max | 1041109,00 | 643959,00 | 397150,00 | 0,73 | 0,53 |

**Table 4.** Chain proportions for each mouse project.

| bioproject | organism | igh | igk | igl | igk_ratio | igl_ratio |
| --- | --- | --- | --- | --- | --- | --- |
| PRJNA980254 | mouse | 2109 | 1942 | 167 | 0.92 | 0.08 |
| PRJNA921879 | mouse | 3649 | 3466 | 183 | 0.95 | 0.05 |
| PRJNA844903 | mouse | 1694 | 1653 | 41 | 0.98 | 0.02 |
| PRJEB49019 | mouse | 173847 | 161748 | 12099 | 0.93 | 0.07 |
| PRJNA845790 | mouse | 4735 | 4496 | 239 | 0.95 | 0.05 |
| PRJEB34451 | mouse | 6883 | 6679 | 204 | 0.97 | 0.03 |
| PRJNA777966 | mouse | 70770 | 66843 | 3927 | 0.94 | 0.06 |
| PRJNA990769 | mouse | 3161 | 2700 | 461 | 0.85 | 0.15 |
| PRJNA1082281 | mouse | 81818 | 54478 | 27340 | 0.67 | 0.33 |
| PRJNA934380 | mouse | 15284 | 14710 | 574 | 0.96 | 0.04 |
| PRJEB64695 | mouse | 144667 | 131659 | 13008 | 0.91 | 0.09 |
| PRJNA1026914 | mouse | 1427 | 1388 | 39 | 0.97 | 0.03 |
| PRJEB50082 | mouse | 12044 | 11182 | 862 | 0.93 | 0.07 |
| PRJNA839019 | mouse | 111798 | 107354 | 4444 | 0.96 | 0.04 |

**Table 5.** Grouped metrics of chain proportions of sequences from mouse projects.

| metric | igh | igk | igl | igk_ratio | igl_ratio |
| --- | --- | --- | --- | --- | --- |
| mean | 45277,00 | 40735,00 | 4542,00 | 0,92 | 0,08 |
| std | 60213,00 | 55233,00 | 7897,77 | 0,08 | 0,08 |
| min | 1427,00 | 1388,00 | 39,00 | 0,67 | 0,02 |
| max | 173847,00 | 161748,00 | 27340,00 | 0,98 | 0,33 |

As described by Collins and Jackson (Collins and Jackson, 2018), the proportion of kappa to lambda light chains varies between the species and might be dependent on the organism’s size. While the low lambda usage in rodents could be partly due to limited combinatorial diversity as only four possible rearrangements of lambda V and J genes, it is not the only reason. Germline usage frequencies are highly variable and dependent on factors such as gene accessibility and their positions within the genome. Moreover, secondary rearrangements reinforce kappa gene usage.

We manually curated metadata for each bioproject, focusing on human and mouse datasets (Figure 2). Human sequences make up approximately 90% of the dataset (Figure 2A), reflecting their prevalence in antibody studies. Using the RIOT program, we assigned constant region (C gene) allotypes to the antibody sequences by aligning their constant regions to known human genes. For heavy chains, the majority were assigned as IGHM, followed by IGHG1 and IGHA1, with about one million sequences remaining unassigned (Figure 2B). For light chains, most sequences were assigned as IGKC, highlighting the prevalence of kappa light chains, while lambda chain assignments were split among IGLC2, IGLC1, and IGLC3; approximately 0.5 million sequences were unassigned (Figure 2C).

**Figure 2.**
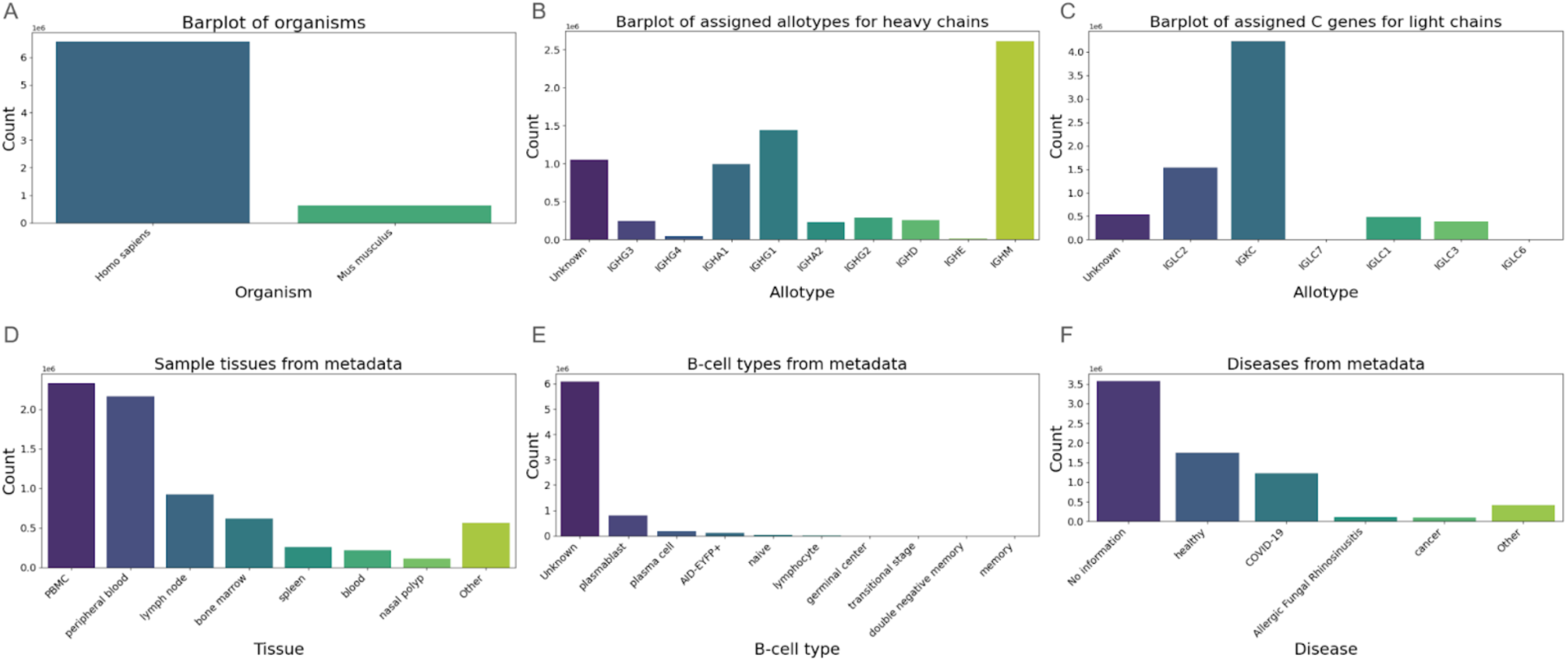
Metadata associated with the 58 paired heavy/light bioprojects. **A.** Sequences count stratified by organisms. **B.** Heavy chain allotypes assigned by C region alignment. **C.** C genes count assigned by C region alignment. **E.** Sequences stratified by source tissue. E. B cell types extracted from metadata. F. Associated diseases

Analyzing tissue sources, we found that the largest number of sequences originated from peripheral blood mononuclear cells (PBMCs), closely followed by peripheral blood samples (Figure 2D). Significant numbers also came from lymph nodes (about 1 million sequences), bone marrow (0.7 million), and the spleen (0.3 million), with other tissues contributing roughly 0.6 million sequences. Regarding B-cell types, most sequences (roughly 6 millions) were unassigned; among the assigned, plasmablasts constituted the largest group, followed by plasma cells, reflecting the diversity of tissue sources and potential challenges in accurate cell type annotation (Figure 2E). Lastly, when examining disease associations, the majority of sequences were unassigned; among the assigned sequences, those from healthy individuals were most prevalent, followed by sequences from COVID-19 patients, with smaller contributions from cancer patients and cases of allergic fungal rhinosinusitis (Figure 2F).

High-level exploration of our data indicates that it is diverse in terms of sequence counts per bioprojects and it does not have debilitating biases in terms of metadata associated with it. Using the opportunity of such a large dataset, we conducted an exploratory analysis of the pairing preferences. We focused on two aspects that were explored previously, germline-level and residue-level heavy-light pairing preferences.

### Estimating the total number of antibody pairs

Given our large paired antibody sample, we aimed to estimate the total number of paired antibody sequences, similar to the way this is done to estimate the number of species on the basis of a smaller sample. This was previously done for the unpaired datasets by Briney et al. (Briney *et al*., 2019) using the Chao2 estimator (Gotelli and Colwell, 2011). Briefly, the Chao2 estimator is a non-parametric method commonly used in ecology to estimate species richness, particularly effective for datasets where many species are rare or unobserved. It accounts for unseen diversity by considering the frequency of rare occurrences—in our case, antibodies observed only once or twice.

Using the Chao2 estimator on our paired antibody sequence database, with the observed number of unique heavy-light paired sequences to be 339,192 (S_observed_), the number of sequences occurring only once equal to 3,390,785 (q_1_) and the number of sequences occurring twice equal to 1139 (q_2_), we calculated the possible number of unique antibodies to be around 5 billion using the equation 1.

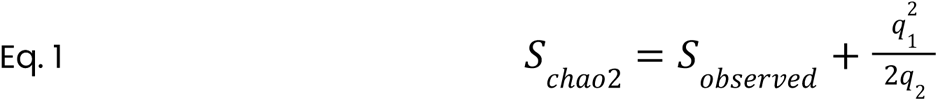

On one hand, this appears to be a gross underestimate, given the known theoretical diversity of the variable regions. While this estimate highlights significant diversity within our dataset, it is lower than some other sources that suggest a higher magnitude of antibody variability of 10^16^ to 10^18^ (Briney *et al*., 2019). This underestimation could be a lower bound estimate of the richness as it has been reported for Chao2 (Chiu, 2023). Moreover, a small sampling ratio could influence the estimated results.

On the other hand, it is known that though there is great theoretical variability of the antibody repertoires, there is evidence that the vast sequence space might be constrained in the structural and sampling fashion (Rees, 2020; Dudzic, Chomicz, *et al*., 2024; Krawczyk *et al*., 2019, 2018). In particular, the variable regions of natural antibodies secreted by B-cells in the human body need to be significantly charged to assure their solubility, stability, and long circulation times (Reddy *et al*., 2010; Lee *et al*., 2019). Furthermore, constant regions of the monoclonal antibodies are highly charged and likewise charged variable regions help prevent antibody aggregation (Li *et al*., 2014). This could be a potential explanation of how such a vast sequence space for antibodies is efficiently restricted by our immune system to a smaller functional repertoire. As pointed out by Collins and Watson (Collins and Watson, 2018), the human light chain repertoire exhibits a surprising lack of diversity due to several constraining factors. A primary reason is the strong bias in light chain gene usage. In the kappa light chain (IGK), just six IGKV sequences dominate reported rearrangements. Similarly, in the lambda light chain (IGLV), only three IGLV genes account for over 50% of human rearrangements, and the utilization of the four functional human IGLJ genes is notably biased. Additionally, the absence of D genes in light chain rearrangements further limits diversity.

With more paired sequences that appear to be in a steady supply (Figure 1), we will be able to see whether the diversity reaches a plateau. At this stage, we would interpret the results from Chao2 as informative but not conclusive.

### Germline encoded pairing preferences

In humans and mice, when the initial rearrangement of the kappa light chain gene is unproductive or generates a self-reactive B-cell receptor, the B-cell can engage in additional rounds of secondary rearrangement through a process called receptor editing (Collins and Watson, 2018). This mechanism enables the B-cell to modify its receptor specificity by replacing the problematic light chain, thereby resolving auto-reactivity. We investigated the presence of pairing preferences between the variable (V) germline genes of heavy and light chains, analyzing kappa and lambda chains separately. Germline usage stems from its accessibility and position in the genome. While some chain pairings arise due to germline usage frequencies in the dataset, we expect that receptor editing introduces a bias. Consequently, we anticipate that highly prevalent heavy-light chain pairs will exhibit lower auto-reactivity.

Firstly, we performed a Chi-squared test on the contingency table for the entire dataset, where columns and rows represented heavy and light V genes respectively. This test confirmed significant pairing preferences in the dataset for both kappa and lambda chains (p << 0.01). To further investigate such preferences, we constructed contingency tables for each pair of heavy-light V gene subgroups. The null hypothesis was that H/L pair frequency is the same as the frequencies of H and L in the whole dataset (see methods section). A detailed description can be found in the methods section. This approach allowed us to assess pairing preferences at the V gene subgroup level by comparing their distributions to the remaining pairs in the dataset, following methodologies from previous studies (Jayaram *et al*., 2012). The results are displayed as heatmaps of counts for kappa and lambda chains (Figure 3) as well as heatmaps for p-values representing the likelihood of pairing frequencies being a consequence of germline usage frequencies (Figure 4). Because we are looking at individual pairs, we applied Bonferroni correction, however the very low p-values still show statistical significance.

**Figure 3.**
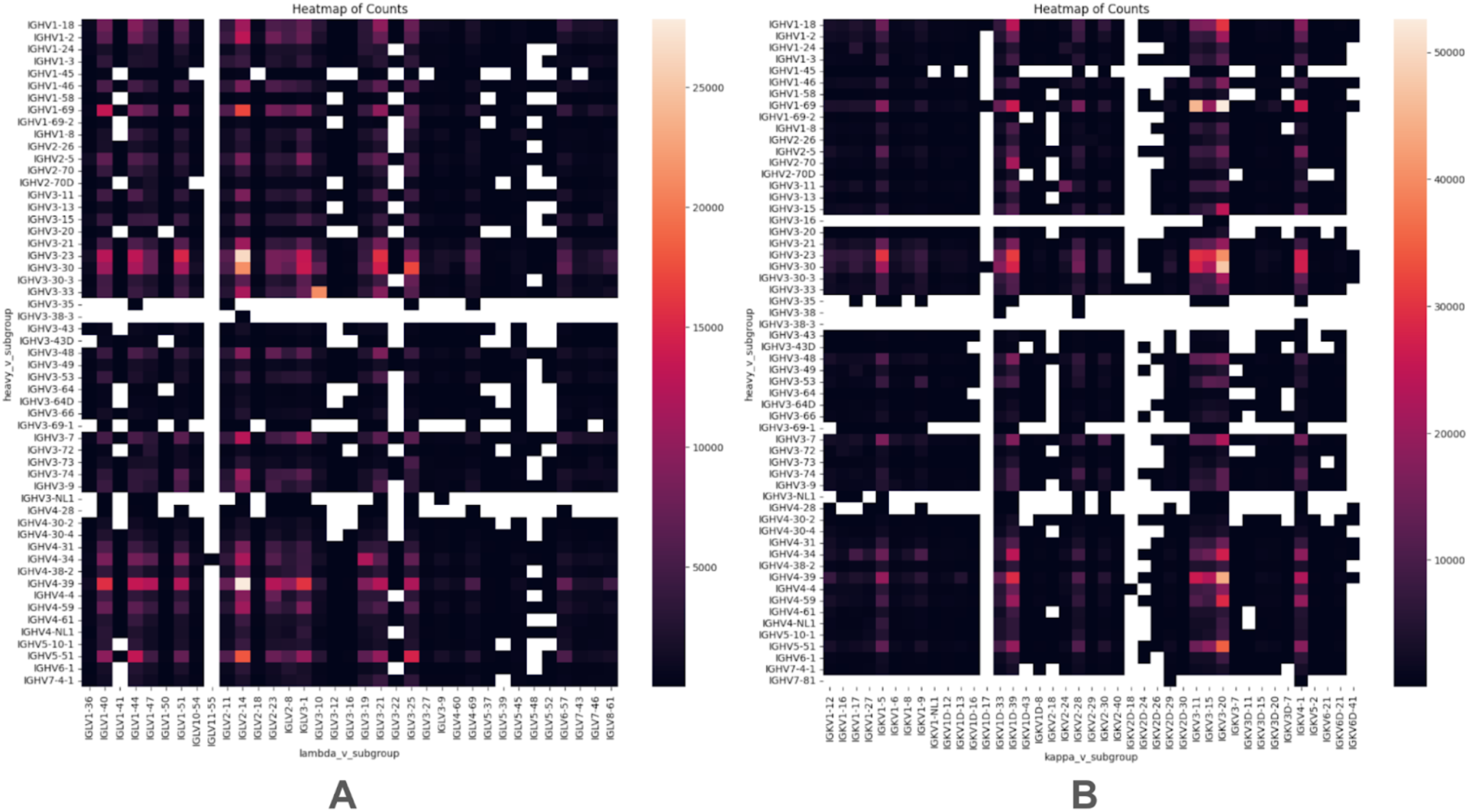
V gene subgroup pairing counts. **A.** Lambda chains. **B.** Kappa chains. Heatmaps represent observed counts of V gene subgroup pairs in heavy and light chains.

**Figure 4.**
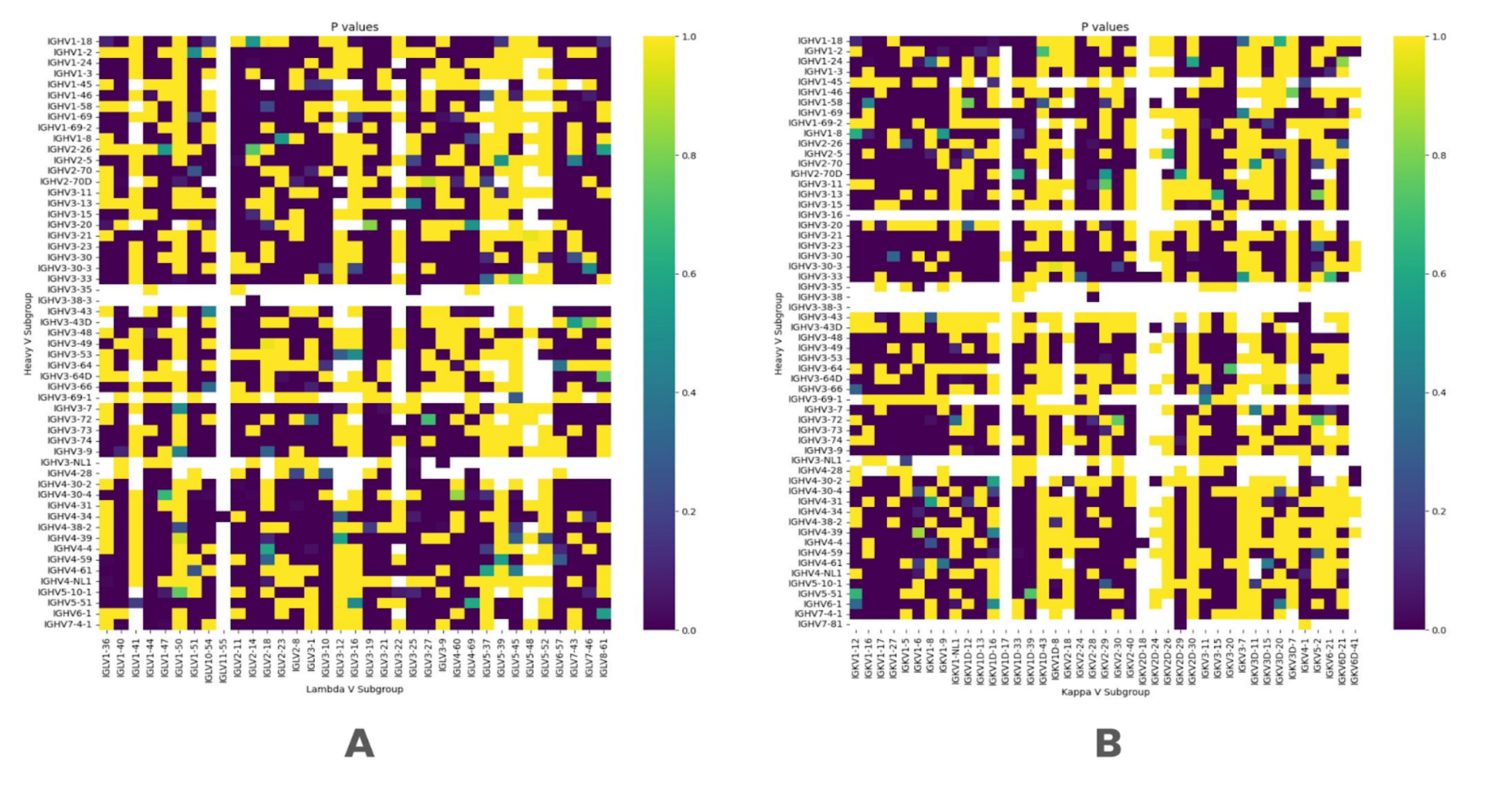
V gene subgroup pairing preferences. A. Lambda chains. B. Kappa chains. Heatmaps represent p-values for individual pairs, indicating how likely we would be to observe a given pair count given the marginals of gene counts in the dataset.

Altogether, we note that there are visible biases in the counts of certain pairs (Figure 3), and there exists a mixed picture of pairs that exist beyond random likelihood, and those that appear purely by chance. These findings indicate that while some pairings result from gene distribution patterns, certain pairing preferences do indeed exist.

To evaluate whether our database provides a unique and enriched perspective on germline gene usage compared to existing public repositories, we conducted a comparative analysis with the OAS paired database. Employing the chi-square contingency test, we tested the null hypothesis that the germline assignment ratios are identical between the datasets. The analysis yielded a chi-square statistic of 138,899 with 55 degrees of freedom for the human heavy chain V genes, resulting in a P-value much smaller than 0.001. Similarly, comparing kappa and lambda chains yielded p-values close to zero, with chi-square of 19,965 and 17,588 and with 33 and 39 degrees of freedom respectively. This highly significant result indicates that the proportions of germline gene usage differ substantially between the two datasets.

Consequently, our database significantly enriches the diversity of germline genes represented, offering a broader and more comprehensive perspective of antibody germline usage than currently available in other databases. Following the Chao2 estimator findings, it is also likely true that our current dataset has intrinsic biases, and as more paired antibody datasets come in, the natural germline pairing distributions among human and murine antibodies may be refined further to contain fewer biases.

### How naturally-sourced pairs compare to clinical-stage monoclonal antibodies

Comparing naturally sourced datasets such as OAS and PairedAbNGS, assumes an intrinsically similar, natural, sampling distribution. This is unlikely the case for monoclonal antibodies, where one can expect that the pairing preferences reflect engineering practices rather than natural preferences. While the lead sequence optimization does not involve disturbing the light and heavy chain pairing of the parent clone for the fear of losing potency, some germlines are preferred due to their favorable liability profiles (Satława *et al*., 2024).

To assess whether germline preferences in monoclonal antibodies align with those observed in natural repertoires, we plotted a heatmap that counted the usages of heavy and light chain genes in monoclonal antibodies (Figure 5). This analysis uncovered a (known) strong bias in monoclonal antibody design toward the IGHV3-23 and IGHV1-46 germlines. The IGH3-23 contains a motif in the V-domain that is capable of protein A binding (Graille *et al*., 2000). This preference is likely attributed to its prominence in observed repertoires and its favorable liability profile (Satława *et al*., 2024), making it a popular choice in therapeutic antibody development.

**Figure 5.**
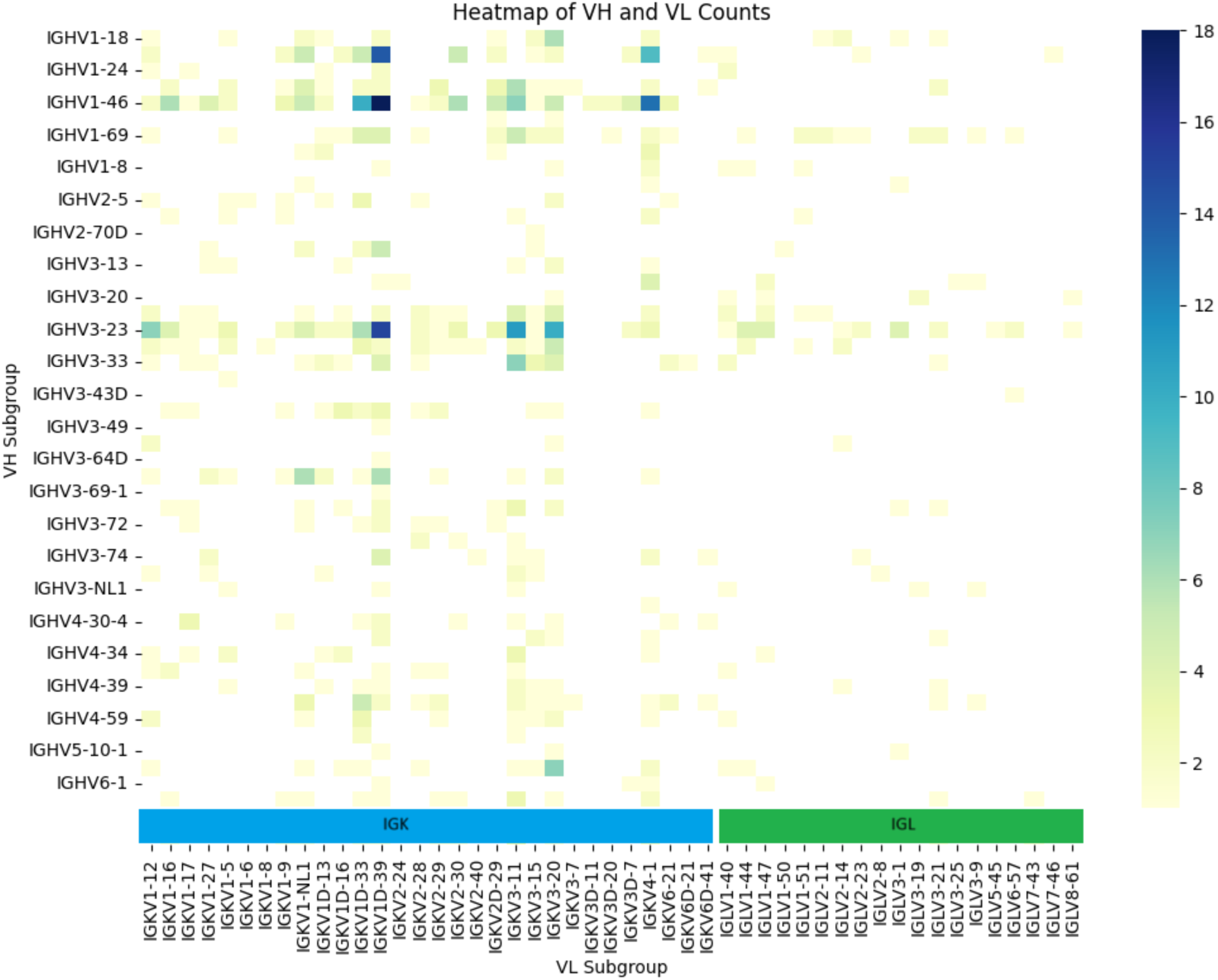
Heatmap of V gene subgroups in monoclonal antibodies. The counts represents the number of antibodies sharing the gene pair.

Another evident bias is the preference for Kappa light chains (Figure 5). This is expected, as many therapeutics are derived from murine parental clones, via humanization, where Kappa chains constitute about 95% of the light chains. While Lambda chains have been reported to have poorer developability profiles because of higher average hydrophobicity (Raybould *et al*., 2024), this is a misconception. Only VL3 germline antibodies contain an aggregation prone region (APR) in the light chain FR2-CDR2 region which leads to aggregation, and can be easily fixed (Nichols *et al*., 2015; Kumar *et al*., 2018). Furthermore, antibodies containing Kappa chains may be more prone to self-reactivity as it was suggested by Hedda Wardemann (Wardemann *et al*., 2004). Moreover, Lambda chains may provide stability during an ongoing immune response, as their codon usage reduces the likelihood of structural changes arising from accumulating somatic point mutations (Collins and Watson, 2018).

Overlooking lambda light chains in therapeutic antibody development can be counter-productive as it limits our ability to find potential binders against a target in the early stages and therefore the lead molecule choices of the antibody-based treatments. Adapting humanization techniques to include lambda light chains can mitigate the existing bias toward kappa chains in therapeutic antibody development, potentially leading to therapeutics with better efficacy, safety, and stability profiles that are more representative of the human immune response.

### Structural analysis of contact residues on the heavy-light interface

At a finer granularity beyond germline pairing preferences, we explored residue contacts between antibody heavy and light chains as they are essential for understanding the molecular basis of their pairing specificity and overall function (Raybould *et al*., 2021). By analyzing these interactions, we studied key residues that contribute to the stability and conformation of antibodies, which has implications for antigen binding and therapeutic antibody design.

We explored antibody structures, deposited in RCSB to identify contact residues responsible for VH/VL interfaces. A similar study was carried out by Raybould et al using Random Forests (Raybould *et al*., 2021). To explore those contact residues, the key interface residues, we imported antibody structures from the RCSB Protein Data Bank and extracted residues between the chains based on a distance threshold of 4.5 Å between any pair of heavy atoms from the heavy and light chains. We present how often those residues are in contact (Figure 6). Heatmap of these contacts was plotted to visualize interaction patterns (Figure 7), and barplot/histograms for each IMGT position were generated to identify the type and frequency of conserved contact residues for heavy and light chains (Figure 8).

**Figure 6.**
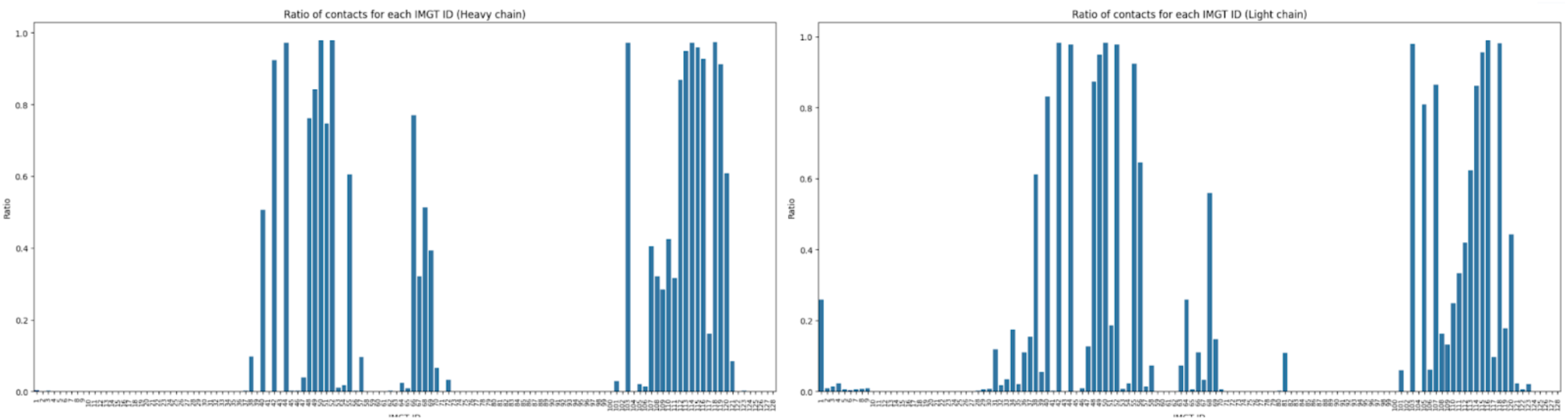
Relative frequencies of residues to be in contact in RCSB Protein Data Bank. For each IMGT ID, we calculated how often its heavy atoms are within 4.5Å of the other antibody chain. Please note that certain IMGT IDs have higher occurrences, and the percentage here is taken as the number of contacts for a given IMGT ID out of the total number of times the residue occurs in the PDB.

**Figure 7.**
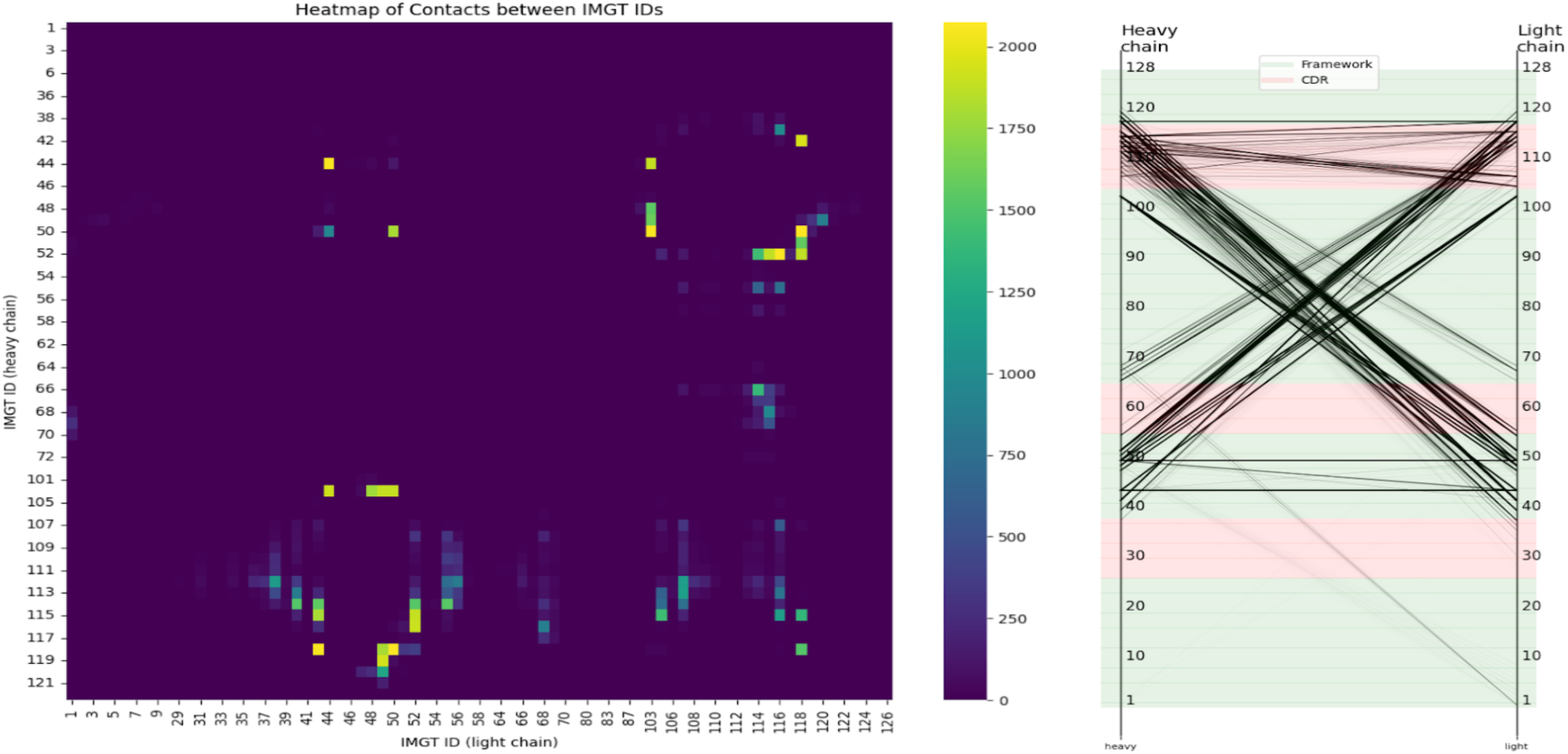
RCSB derived IMGT contact residues for heavy and light chains. Left: heatmap of counts of contacting residues between heavy and light chains as defined by 4.5 Å distance threshold. Right: Slope chart presenting inter-chain contacts.

**Figure 8.**
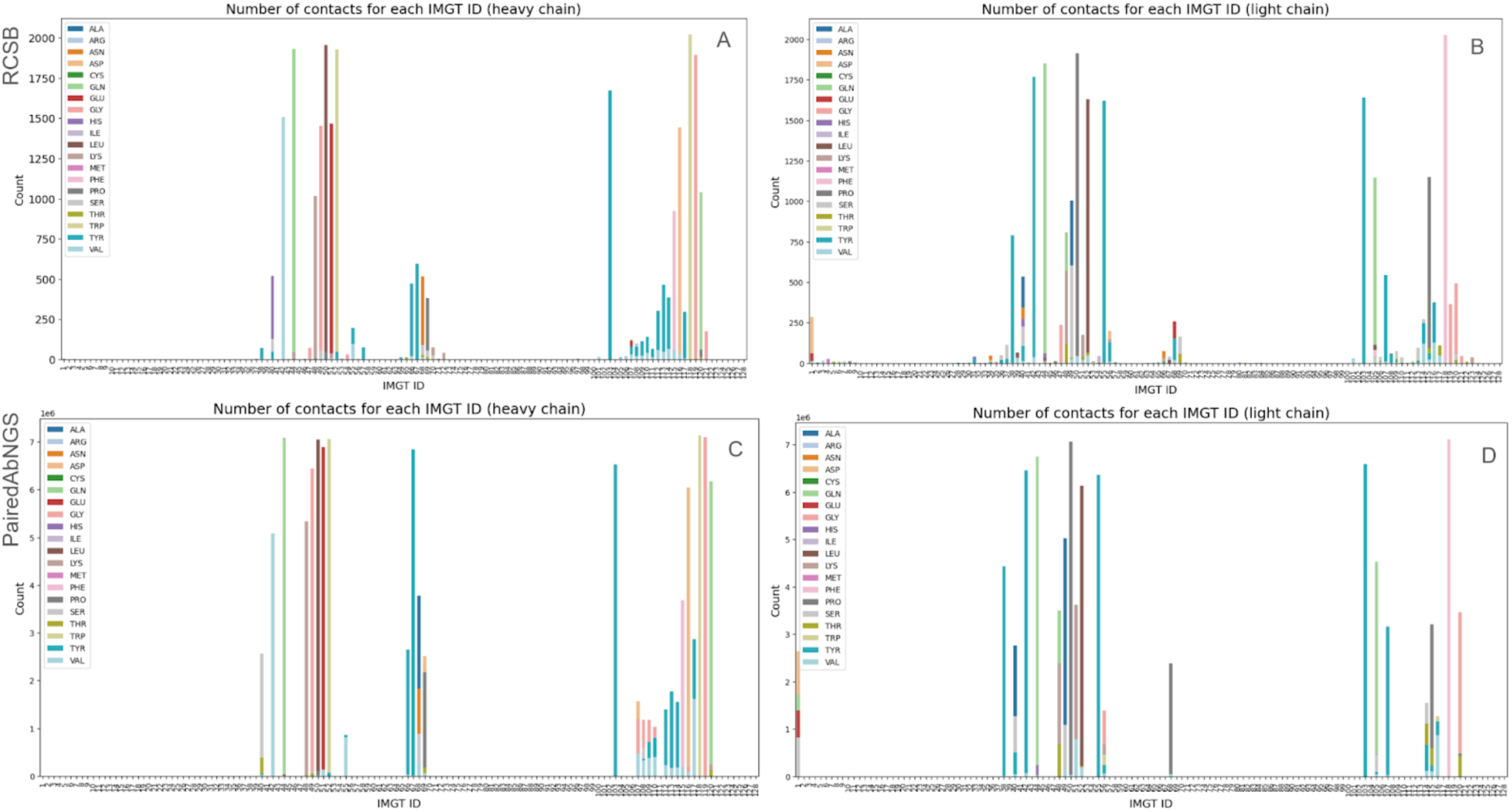
Amino acid distributions for key interface residues. For each key interface residue (<4.5Å heavy atom distance in the PDB), we plot how often the position is seen in a given dataset (y-axis count) and on the bar, we color by proportion of amino acid types at that position. A. Heavy chains from PDB. B. Light chains from RCSB PDB. C. Heavy chains from PairedAbNGS. D. Light chains from PairedAbNGS.

Such proximal residues might affect VH/VL packing and thus affect binding affinity to the target. Notably, we observed that the CDR3 of each chain is in contact with the FWR2 of the opposing chain. This suggests that while the FWR2 region of one chain does not typically interact with antigens, it may influence the conformation of the CDR3 loop from the other, thereby affecting antigen binding. Influence of FW2 on the conformation of CDRs is a well-known issue in humanization. Additionally, the interaction between CDR3 loops contributes to chain pairing specificity, underscoring the importance of these regions in antibody structure and function. Our findings are consistent with previous studies that have reported similar interactions (Abhinandan and Martin, 2010; Boron and Martin, 2023; Fernández-Quintero *et al*., 2020; Cisneros *et al*., 2019; Dunbar *et al*., 2013; Bujotzek *et al*., 2015) using smaller datasets.

We compared our key interface residues to Vernier residues as defined by Foote and Winter (Foote and Winter, 1992) for heavy and light chains (Figure 9). Vernier residues were translated to IMGT using IMGT lookup. We observed very small overlap between the positions.

**Figure 9.**
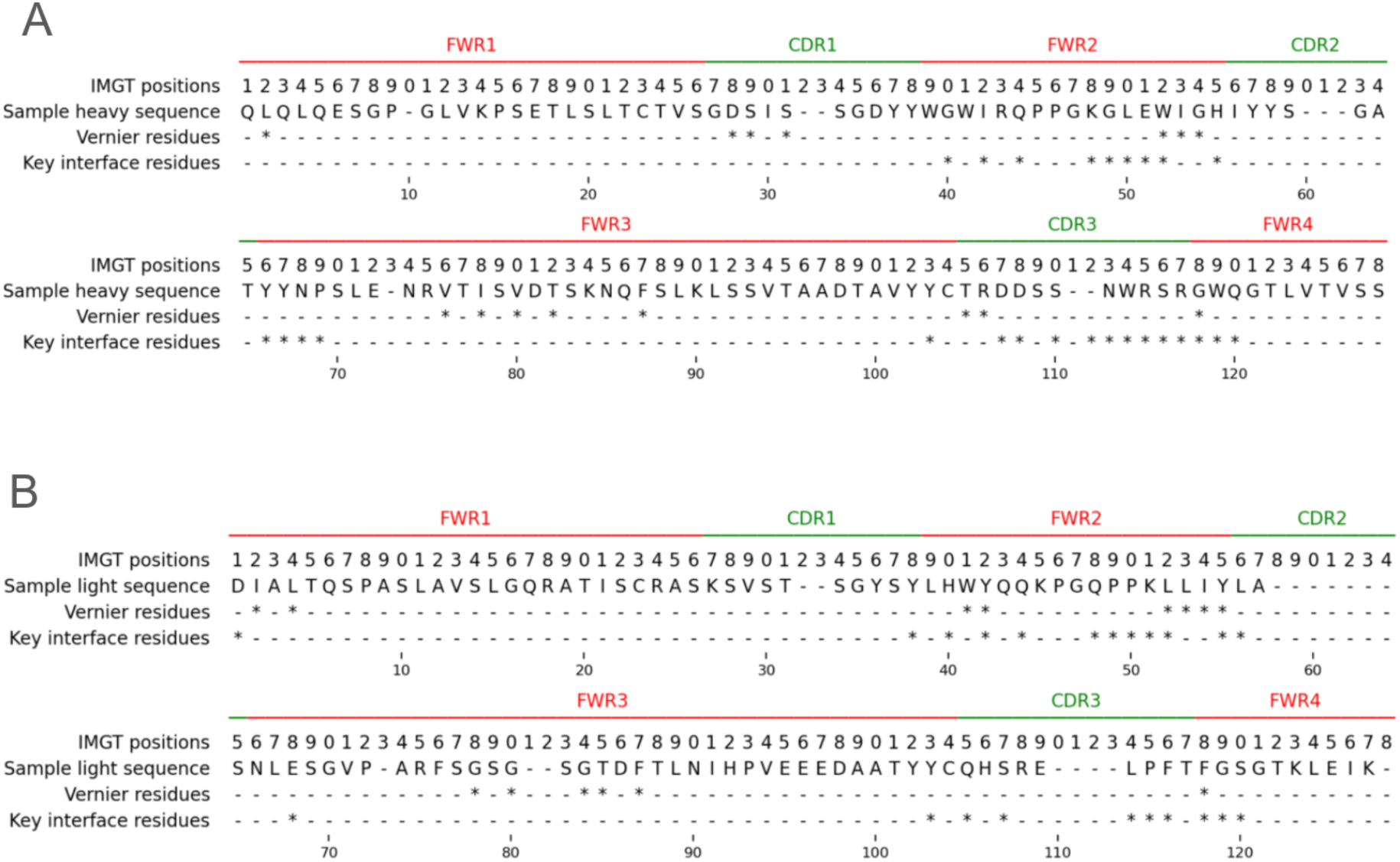
Key interface residues compared to Vernier residues in the IMGT numbering scheme. **A.** Heavy chain. **B.** Light chain.

Building upon the key interface residues extracted from the PDB, we analyzed their amino acid distribution in our large paired sequence database. For each key residue on either heavy or light chain, we plotted the distributions of amino acids on key interface residues (Figure 8 C,D). As the distributions qualitatively appeared similar, we quantified the difference between them using Total Variation Distance, defined as half of the sum of proportions of each amino acid, as defined by equation 2:

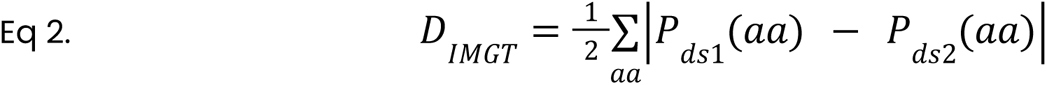

Here, the distance D_IMGT_, calculated for specific IMGT position, measures how the distributions of amino acids are different at that position between the datasets. P gives the proportions of a given amino acid (aa) from a datasets (e.g. PairedAbNGS, PDB). Therefore the sum of all absolute values of differences between the proportions P in the datasets measures the absolute difference between their proportions. Please note the sum of the proportions is one at any position, for any dataset, normalizing for biases of different numbers of certain residues at different positions (Figure 10).

**Figure 10.**
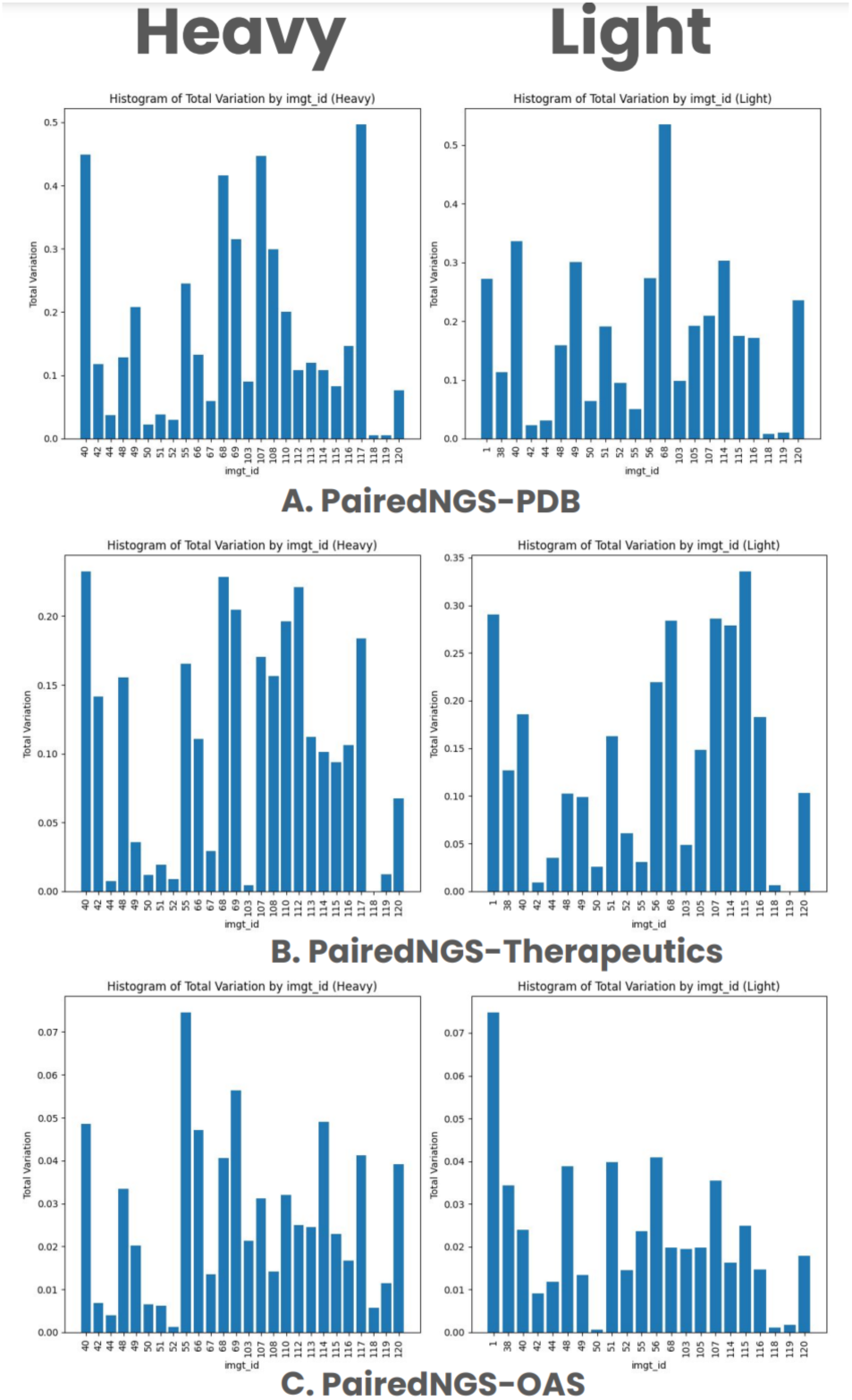
Total Variation Distances for each key residue between amino acid distributions. The closer the value is to zero, the closer the distributions. **A.** PairedAbNGS vs PDB. **B.** PairedAbNGS vs therapeutics. **C.** PairedAbNGS vs OAS.

We calculated such measures between PairedAbNGS and PDB (Figure 10A), therapeutics (Figure 10B), and OAS (Figure 10C). Most of the values are below 0.2, which can be roughly interpreted as 80% similarity between the amino acid distributions. This similarity indicates that the contact residues identified are conserved across both structural, therapeutic, and sequence data, reinforcing their significance in contributing to heavy-light chain pairing specificity. The consistent patterns suggest that these residues play a pivotal role in maintaining the structural integrity and functional conformation of antibodies.

To further examine whether amino acids distributions on key interface residues differ between databases, we performed a series of goodness of fit chi-square tests comparing our database, monoclonal antibodies, Paired OAS, and RCSB PDB. For each pair of databases (PairedAbNGS vs other), for each key residue identified in the RCSB, we constructed contingency tables where columns represented distinct amino acids. The null hypothesis was that the distribution of amino acids on specific positions is the same therefore the expected value for each amino acid was calculated as the proportion of particular AA on this position in our database, multiplied by the number of data points on the same position from the database compared. Categories with less than 5 observed amino acids were binned to the “other” category and the results were Bonferroni corrected. While histograms with Total Variation Distances revealed moderate and high similarities in some residues distributions, in the majority of cases the null hypothesis was rejected with p∼0 (with the exception of highly conserved residues).

In particular, when comparing key residues between OAS and PairedAbNGS, there were no IMGT positions that showed the same distributions according to Chi square tests. This means that while highly similar (Figure 10), PairedAbNGS extends the diversity landscape of the available data, not only in germline gene usage but also in sequence variability. The observed difference underscores that PairedAbNGS data can reshape our understanding of antibody diversity, with downstream implications for modeling, engineering, and applying immunoglobulin repertoires in both research and clinical contexts.

We investigated the inter-chain amino acid pairs of the previously identified pairs of key interface residues to gain deeper insights into their contributions to heavy-light chain pairing specificity. For each pair of IMGT positions corresponding to these key residues on the heavy and light chains, we calculated the Total Variation Distance of amino acid pairs. The Total Variation Distance served as a measure of the difference between the distribution of amino acid pairs in our dataset and the distribution expected if heavy and light chains were randomly paired. By plotting these results as a heatmap (Figure 11), we visualized the extent of these differences across all residue pairs. Generally, the differences were small, likely due to random pairings being a very poor proxy for ‘incorrect pairing’ as they occasionally replicated correct pairings present in the actual dataset. However, we observed notable differences reaching up to 0.06 in Total Variation Distance values, indicating that the observed amino acid pairings differ significantly from random expectations at certain positions. In particular, the majority of significant differences were attributed to the CDR3 region of the heavy chain, reinforcing its critical role in chain pairing specificity. This finding suggests that specific amino acid interactions involving the heavy chain CDR3 region are key determinants in the proper pairing of antibody chains. This aligns with work of Monica L. Fernández-Quintero et al (Fernández-Quintero *et al*., 2022) who found that specific residues within the CDR-H3 and CDR-L3 loops can determine pairing preferences.

**Figure 11.**
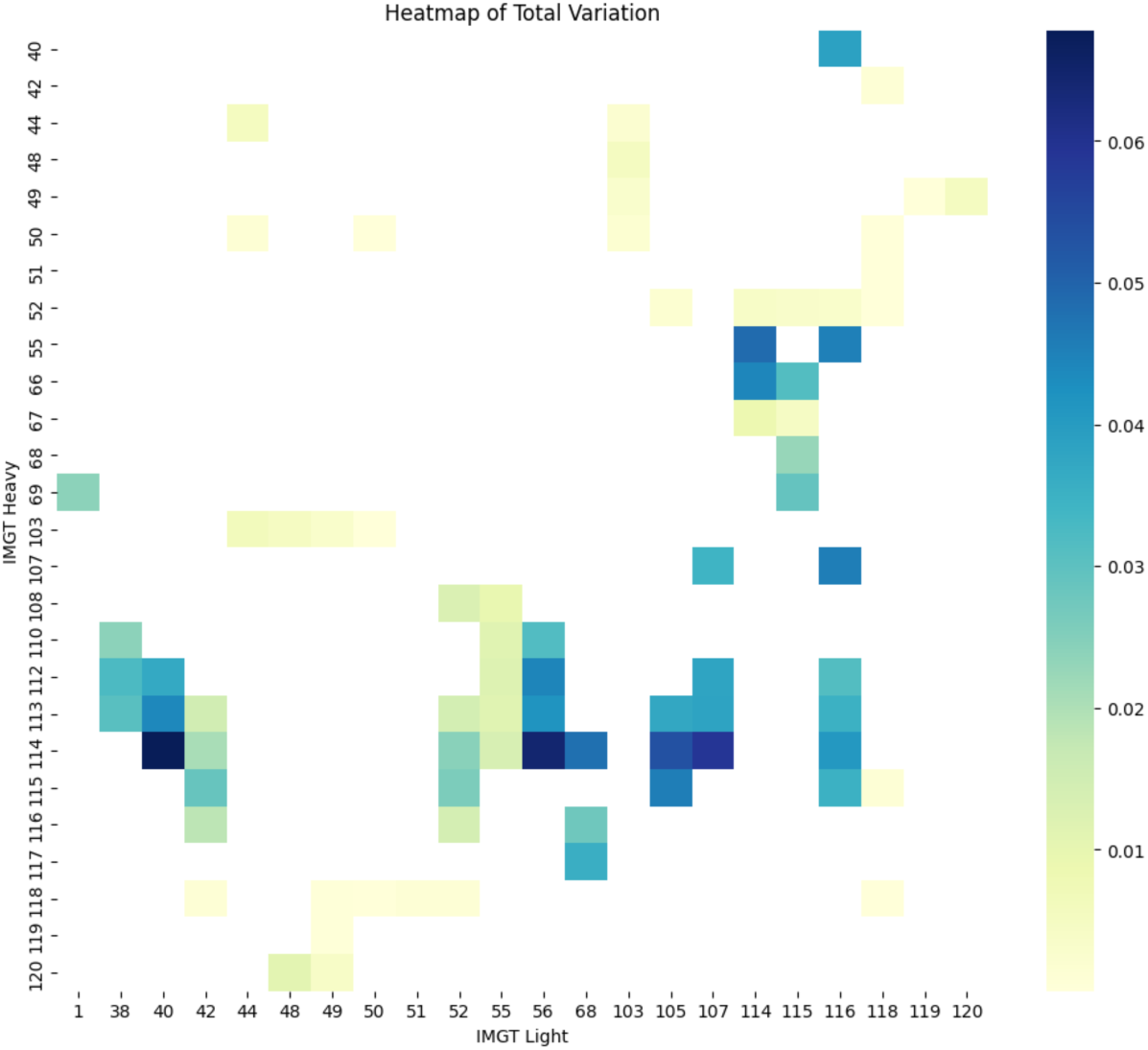
Total Variation Distances for each pair of key residues between amino acid distributions.

Having identified the proximal contact of CDR3 and FWR2 from the pairing chains (Figure 7), we explored the pairing preferences of inter-chain V and J genes (Figure 12). Similar to Figure 3, we observe a strong bias towards some pairings (p-value ∼ 0) which also might indicate pairing preferences arising from receptor editing in which autoreactive antibodies are removed. It is important to note however, that while we observe a notable bias in J gene usage towards IGHJ4, it can also be attributed to recombination signal sequence (RSS) as it was investigated by Bin Shi et al (Shi *et al*., 2020).

**Figure 12.**
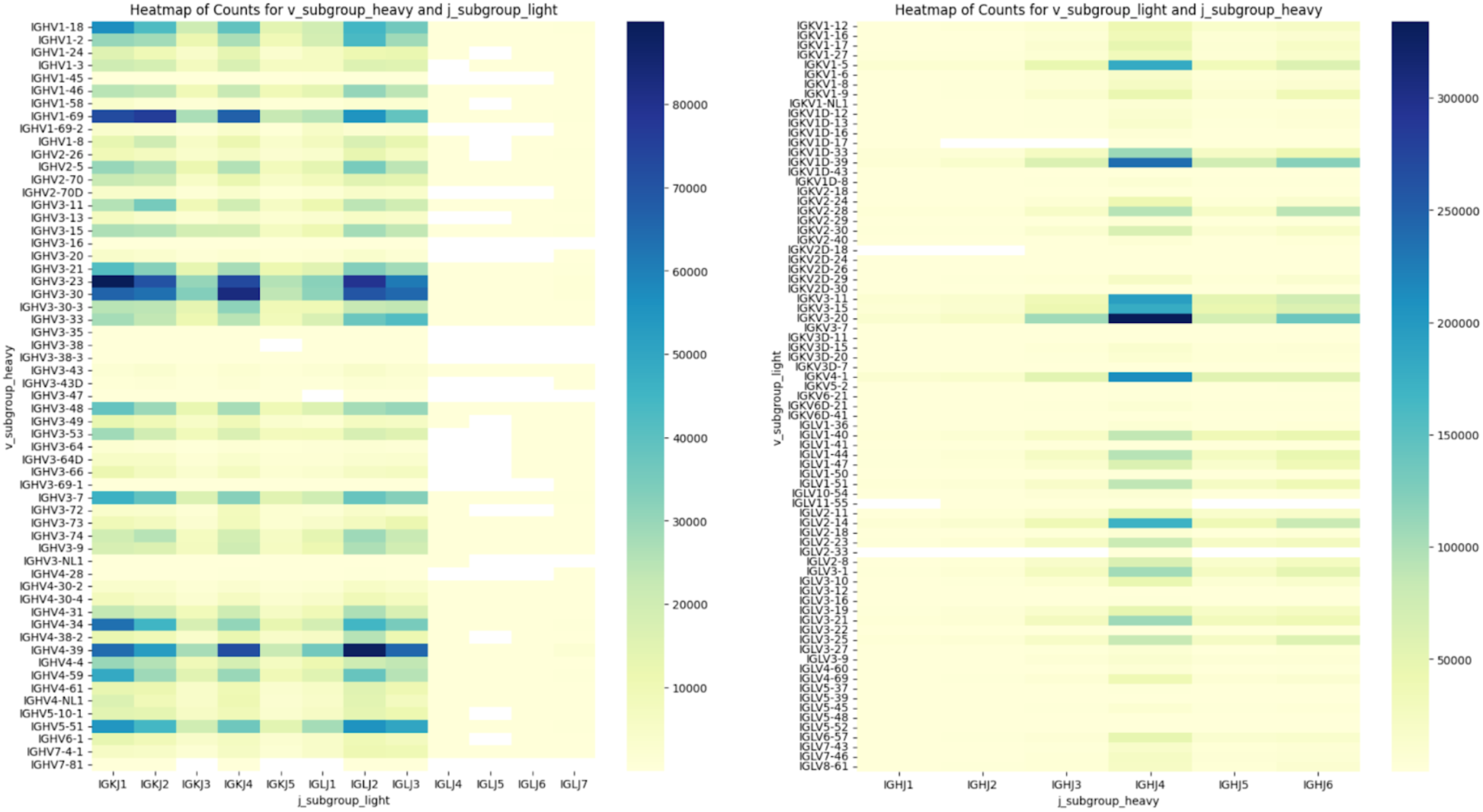
Inter-chain V-J germline pairing preferences.

## Discussion

We introduce PairedAbNGS, which to our knowledge is the largest available dataset of paired heavy and light chain antibody sequences. This extensive dataset serves as a valuable resource to the community for several applications including repertoire analyses, biologic drug discovery and machine learning applications in antibody design and engineering. By providing a comprehensive collection of paired sequences, PairedAbNGS enables Machine Learning models to learn the intricate relationships between heavy and light chains, leading to more accurate functional antibody predictions and the development of more effective antibody libraries with higher sensitivity and specificity along with desirable physicochemical attributes.

We examined pairing preferences between the variable (V) genes of heavy and light chains and identified significant biases toward certain pairs that go beyond gene usage frequencies alone. Using Chi-square contingency tests, we confirmed that specific heavy-light chain combinations are favored, likely due to receptor editing favoring combinations of stable molecules, with lower autoreactivity.

Exploring the key regions and residues at the heavy and light chain interfaces, we noted that the framework region 2 (FWR2) from one chain frequently contacts the complementarity-determining region 3 (CDR3) of the pairing chain. This observation led us to investigate pairing preferences for VH-JL (heavy chain variable region and light chain joining region) and VL-JH (light chain variable region and heavy chain joining region) gene segments. We identified statistically significant biases in these pairings, indicating that specific inter-chain V and J gene combinations contribute to stability and functionality of the antibody molecule.

These biases have important implications for antibody engineering, particularly in the humanization of monoclonal antibodies and the better utilization of lambda chains, which are reported to have lower autoreactivity. Heavy and light chain pairing is crucial for maintaining the correct antibody structure and function. Therefore during humanization, it is essential to consider not only the CDRs but also these key interface residues that contribute to chain pairing specificity and stability. Understanding these pairing preferences and inter-chain interactions can inform strategies to minimize autoreactivity and improve the efficacy of therapeutic antibodies. Our findings highlight the importance of considering heavy-light chain pairing biases and interfacial residues in antibody design and engineering.

As we focused in our analysis on static structures deposited in RCSB, it is important to notice that CDR3 loops are flexible and their lengths are dependent on Vh, Vl and Jh gene usage (Sankar *et al*., 2020). It is therefore important that the pairing preferences, the nature of inter-chain contacts, and its impact on antigen recognition should be investigated using conformational ensembles obtained from molecular dynamics simulations as it was introduced by Monica L Fernández-Quintero et al. The authors showed that varying light and heavy pairings impacts CDR conformations. This leads to changes in heavy/light chain orientations which further impacts antibody structure (Fernández-Quintero, Georges, *et al*., 2021; Fernández-Quintero, Kroell, *et al*., 2021).

Considering the potential of transcriptomic analysis as a future extension of our database could significantly enhance our understanding of antibody development and maturation. Some of the bioprojects within our database contain sequencing experiments that allow for the determination of protein expression levels. Integrating transcriptomic data would enable us to identify biomarkers relevant to antibody maturation and immune responses, which could be instrumental in discovering antigen-specific antibodies directly. Although we did not incorporate this analysis in the current study due to incompatibilities among data sources, future inclusion of transcriptomic information could provide deeper insights into the regulatory mechanisms governing antibody expression. This expansion would not only enrich the database but also offer a valuable resource for researchers aiming to explore the interplay between antibody structure, expression levels, and immune functionality.

## Methods

### Identification of bioprojects

NCBI SRA (Leinonen *et al*., 2011) was used as the primary source of the sequences. Metadata for all bioprojects was imported from European Bioinformatics Institute European Nucleotide Archive (EBI ENA) portal (Cummins *et al*., 2022).

We extracted sequencing runs from NCBI SRA metadata where the library source was defined as “TRANSCRIPTOMIC SINGLE CELL”. For each unique bioproject we imported its description from EBI ENA. In order to identify antibody-related projects, we constructed an LLM prompt and submitted it to the Gemma language model. Finally, the remaining projects were manually reviewed to discard studies incorrectly flagged as antibody-related. Furthermore, we applied the following inclusion criteria: 10x Chromium 5’ library, human or mouse, known data layout.

Prompt engineering (White *et al*., 2023) is an evolving field focused on optimizing interactions with large language models to achieve specific outcomes. It involves crafting precise prompts to guide LLMs in generating accurate and context-appropriate responses, utilizing techniques such as chain of thought and emotion prompts to enhance problem-solving and emotional intelligence. Additionally, few-shot learning and self-reflection prompts are used to help LLMs quickly adapt to new tasks and improve the quality of their outputs. We have applied the above-mentioned prompt engineering patterns to create the prompt given in Listing 1.

**Listing 1.**
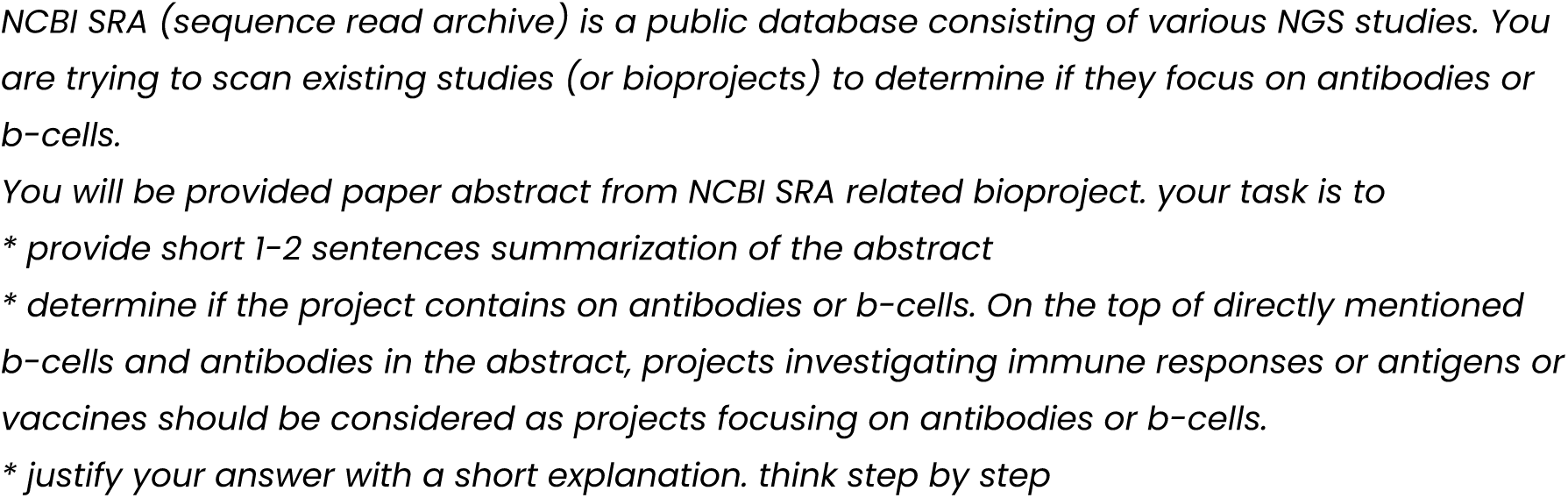
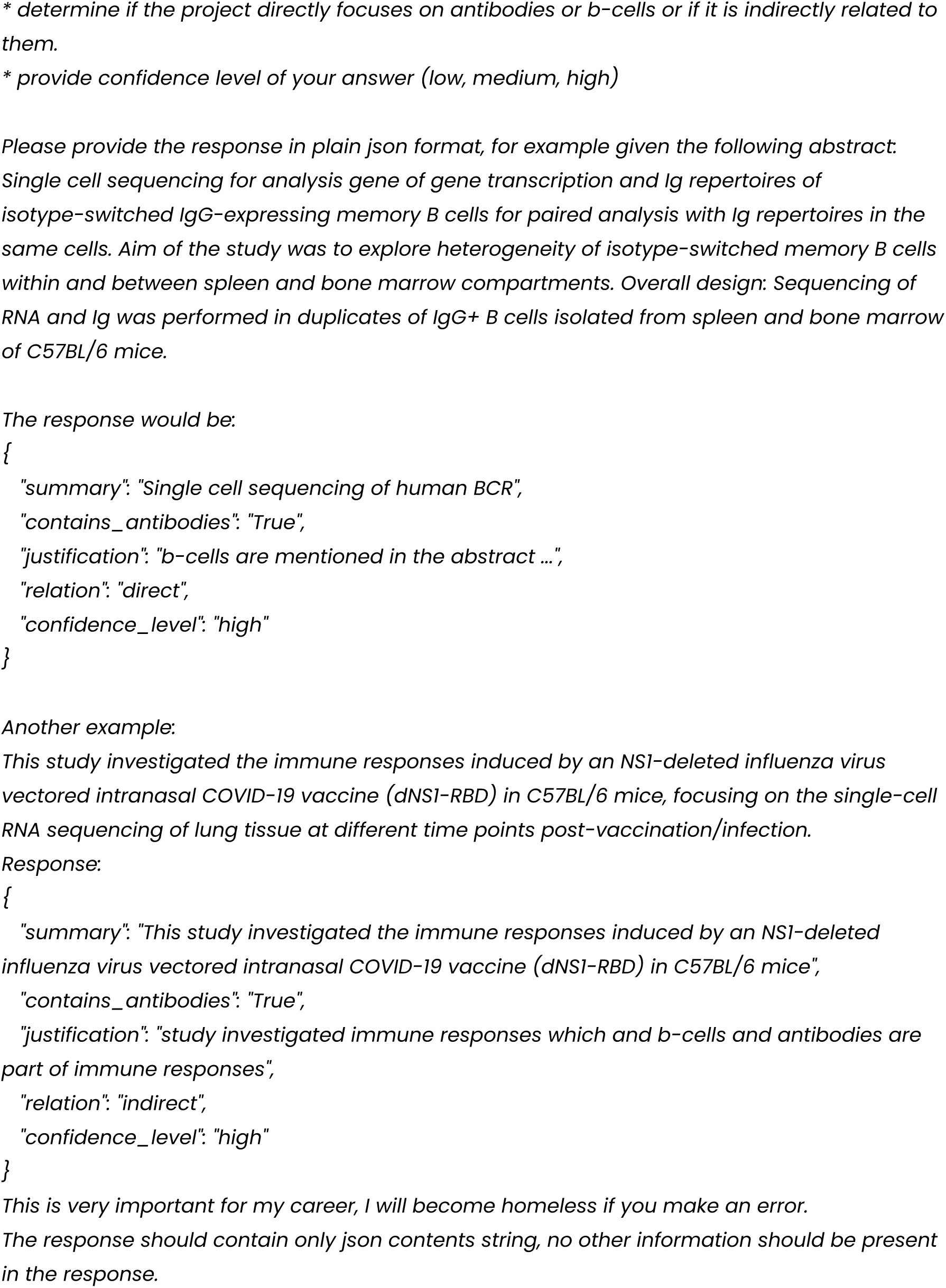
Bioproject verification prompt.

This filtered the results to ca. 200 antibody-related bioprojects. Such a number was already manageable for further manual review that discarded false positives as well as cases that were not suitable for downstream analysis (e.g. species with projects for which we do not have a germline database). We focused on projects where a 10x Chromium 5’ library preparation pipeline was used.

This way we identified 58 studies, consisting of 2482 sequencing experiments with a total over 14 million assembled productive sequences constituting more than 7 million paired heavy and light sequences.

Reads were converted to FASTQ format using sra-toolkit after which were extracted into separate files and assembled using TRUST4. Finally assembled consensus sequences were annotated using RIOT to IMGT scheme. The OGRDB database was used as a primary source for human and mouse VDJ genes in the RIOT annotation pipeline.

### Processing Pipeline

For each identified project, each read run is extracted to multiple fastq files using the sra-toolkit.

We use TRUST4 script to extract the reads to separate files containing barcodes, UMIs, and biological reads respectively. This process is customized for each study, according to the read layout imported from NCBI trace server or library construction protocol described in study related paper (if present). From the existing bioprojects, we identified 2 groups of read layouts: single - where biological sequence was present on only one read - and paired.

In a single layout, read 1 contains the cell barcode and unique molecular identifier (UMI), which are crucial for identifying the cell of origin and removing PCR duplicates, respectively. Read 2 captures the actual cDNA sequence, which corresponds to the transcribed RNA from a particular gene (Cohen *et al*., 2023).

In paired layout, the length of read 1 is extended to cover both the cell barcode and a portion of the transcript sequence, a modification referred to as “10X 5’ with extended R1” (Cohen *et al*., 2023). This approach can increase the number of unambiguous reads spanning important junctions, such as those found in viral subgenomic mRNAs, potentially improving the accuracy and depth of single-cell transcriptome analysis.

Distinct versions of 10x chemistry have different lengths of 10x barcodes and UMIs. While in single read layout, the length of the UMIs and barcodes is determined by read length, in paired reads we inferred it by localizing conserved Template Switch Oligo (TSO) on read 1.

Extracted reads are repartitioned by barcode to multiple directories to improve the parallelisation of the calculations and TRUST4 is used again to assemble consensus sequences for each barcode. Finally, consensus sequences are annotated using RIOT.

### Germline-encoded pairing preferences

To explore pairing preference between V germlines of heavy and light chains for kappa and lambda chains separately, we grouped sequences by the V gene subgroup, according to germlines assigned by RIOT. Since OGRDB mouse germlines do not have a straightforward group/subgroup/allele hierarchy, we focused only on human projects. We constructed a contingency table containing counts of V gene subgroups frequencies for heavy and light chains which was an input to Chi-square test. Pairs with observed co-occurrence of less than 5 were filtered out.

To investigate pairing preferences at higher granularities, for each pair of *hl*, we constructed the following contingency table:

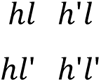

Where *hl* is the count of occurrences of heavy(*h*) and light(*l*) V gene subgroups in the dataset, and *h*’ and *l*’ represents is the number of all other V genes subgroups, except *h* and *l* respectively. The null hypothesis is that the frequency of *hl* stems from the frequencies of *h* and *l* in the dataset, therefore expected values arises from proportion of *hl*, *h*’*l*, *hl*’ and *h*’*l*’. Values were adjusted with Bonferroni correction.

### IMGT key interface residues

We imported protein structures from the RCSB Protein Data Bank and numbered them to the IMGT scheme to identify antibody sequences. To accurately pair heavy and light chains, we measured the alpha carbon (Cα) distances of cysteines located at IMGT position 104 in both chains, pairing those within the distance of 20Å, due to their conserved disulfide bonds. We extracted contact residues between the chains based on a criterion where any pair of heavy atoms (excluding hydrogens) from the heavy and light chains were within a maximum distance of 4.5 Å.

### OAS database preprocessing

The OAS dataset was preprocessed using RIOT to ensure consistent and comparable germline V gene assignments across both databases. We grouped the assigned V genes into their respective subgroups and constructed a contingency table where each row represented a V gene subgroup and each column represented one of the two datasets.

### Therapeutic database preprocessing

Manually collected therapeutic data was filtered to retain only standard monoclonal antibody formats therapeutics (monospecific antibodies with two pairs of heavy and light sequences). Since many therapeutic molecules originate from rodents with varying levels of humanization, sequences were annotated using RIOT using organism detection mode, so that the closest germline can be assigned, regardless of the level of mouse sequence contents.

## Availability

PairedAbNGS is available from https://naturalantibody.com/paired-ab-ngs for non-commercial use by non-commercial organizations.

## Notes

https://naturalantibody.com/paired-ab-ngs/

### Competing Interest Statement

The authors have declared no competing interest.

### Summary of Updates

The authors updated a database link so it points to correct location

